# Beyond Reward Prediction Errors: Human Striatum Updates Rule Values During Learning

**DOI:** 10.1101/115253

**Authors:** Ian Ballard, Eric M. Miller, Steven T. Piantadosi, Noah Goodman, Samuel M. McClure

## Abstract

Humans naturally group the world into coherent categories defined by membership rules. Rules can be learned implicitly by building stimulus-response associations using reinforcement learning (RL) or by using explicit reasoning. We tested if the striatum, in which activation reliably scales with reward prediction error, would track prediction errors in a task that required explicit rule generation. Using functional magnetic resonance imaging during a categorization task, we show that striatal responses to feedback scale with a “surprise” signal derived from a Bayesian rule-learning model and are inconsistent with RL prediction error. We also find that striatum and caudal inferior frontal sulcus (cIFS) are involved in updating the likelihood of discriminative rules. We conclude that the striatum, in cooperation with the cIFS, is involved in updating the values assigned to categorization rules when people learn using explicit reasoning.

## 1.1 INTRODUCTION

Humans possess a remarkable ability to learn from incomplete information, and rely on multiple strategies to do so. Consider a card game where hearts are rewarded and other cards are not. A simple learning strategy, model-free learning, directly associates stimuli and/or actions with rewards that they predict (Sutton and Barto 1998). This algorithm would efficiently learn that card suits predict different reward values. Now consider a more complex game in which the queen of spades is also rewarded, except when it is paired with all the hearts. A more efficient strategy than model-free learning would be to learn the abstract rules or categories that apply to the cards. This strategy requires a cognitive model of the environment based on explicit rules (Miller and Cohen 2001). A large body of work has mapped the neural circuitry underlying model-free learning as well as the circuitry underlying the execution of well-learned cognitive models (Badre and D'Esposito 2009). However, little is known about how cognitive models are acquired or where the variables required to learn models are represented.

Model-free and cognitive model learning have typically been associated with different neural systems: a mesolimbic striatal system for the former and a lateral cortical system for the latter (Glascher et al. 2010; Daw et al. 2011). In model-free learning, striatal neurons encode the value of different stimuli and actions and communicate values to cortical regions via recurrent loops (Haber & Knutson, 2010). Ascending midbrain dopamine projections carry signed reward prediction errors (Montague et al. 1996; Schultz 1997) that underlie the learning of stimulus-and action-outcome associations (Reynolds et al. 2001; Kawagoe et al. 2004; Daw et al. 2005; Morris et al. 2010). Conversely, in cognitive models, prefrontal neuronal pools represent abstract rules, implement control over behavior, and update rules when appropriate (Buschman et al. 2012). We tested the hypothesis that learning cognitive models depends on parallel striatal-prefrontal neural circuitry to what is known to be involved in model-free reinforcement learning (RL).

This hypothesis may appear straightforward, but it faces several theoretical challenges. First, RL and rule-based learning operate on different information: the former assigns values to stimuli or actions and the latter reasons explicitly over abstract rules, concepts, or structured relationships (Goodman et al. 2008; Glascher et al. 2010; Tenenbaum et al. 2011). We propose that, in addition to encoding stimuli and action values, striatal neurons also encode values of the rules represented in cortex (e.g., the value of “all hearts and the queen of spades”). Second, explicit rule-learning requires a different learning signal than the reward prediction error (RPE) calculated in RL. We propose that the dopamine system is not specialized for conveying RPEs; rather, it encodes update signals derived from new information in a variety of learning contexts.

In order to test this hypothesis, we focused on the robust observation in RL research that striatal activation changes in proportion to RPE (McClure et al. 2003; O'Doherty et al. 2003; Rutledge et al. 2010; Garrison et al. 2013). We tested whether the striatum represented RPEs when subjects are biased to learn by reasoning over explicit rules, rather than by incrementally adjusting stimulus-response relationships. If you are learning a card game where hearts are rewarded unless they are paired with a queen, and for many hands you have seen no queens, then discovering that hearts and queens together fail to deliver a reward is highly surprising and also delivers a negative RPE. If the striatum represents RPE as part of a RL algorithm (Garrison et al. 2013), then the striatum should respond negatively to this unexpected and incorrect outcome. Conversely, if the striatum is involved in updating rule values, then it should respond positively to this surprising outcome.

## MATERIAL & METHODS

### 2.1 Participants

Nineteen participants completed the study (11 female; mean age 21.7 years; SD age 7.3 years). Stanford University’s Institutional Review Board approved study procedures, and all participants provided informed consent. Three were excluded because their accuracy was not significantly better than chance. An additional two participants were excluded for excessive head motion (exceeding 2 mm in any direction), leaving 14 participants for analyses. Although this sample size is small, our analyses are focused on very robust and large effects of error response in the striatum, which have been reliably detected with samples of this size (O'Doherty et al. 2003; Rutledge et al. 2010). However, we are underpowered to detect reliable between-subject brain-behavior correlations, and so we do not test for them (Yarkoni 2009).

### 2.2 Rule Learning Task

We used a task that was designed to bias participants towards an explicit rule-learning strategy, rather than an incremental build-up of stimulus-response contingencies. On each trial, a stimulus was shown that varied on three perceptual dimensions (color: blue or yellow; shape: circle or square; and texture: striped or checkered). Participants assigned stimuli to one of two possible categories, “Dax” or “Bim,” based on perceptual features. Participants were informed that rules linking features to categories changed with each new block of trials. Blocks were separated into clearly demarcated scanning runs to minimize interference between rules. Six different rules were learned in counterbalanced order across blocks: A (e.g., blue stimuli are Bims; yellow are Daxes); *A and B* (e.g., blue square stimuli are Bims); *A or B*; *(A or B) and C*; *(A and B) or C*; and *A xor B* (e.g., all blue or square stimuli, with the exception of blue square stimuli, are Bims). The specific features and category labels that defined each rule were randomly determined at the start of each block. Structurally, the order of trials within a given rule block was identical across participants, enabling direct comparison of performance on a trial-by-trial basis. However, the mapping of stimulus features (e.g. A to the color blue, or the shape square, etc.) was randomized. For complex rules, it is necessarily the case that simpler rules sufficed to explain the data for initial trials, and then discriminating examples subsequently require updating to more complex rules.

Each trial was divided into three phases: cue, response, and feedback. Each phase lasted 2 s and was separated by a random 4 to 6 s delay. During the cue phase, the stimulus to be categorized was shown in the center of the screen. During the response phase, a question mark was shown in the center of the screen, which prompted participants to categorize the stimulus by pressing one of two buttons. During the feedback phase, a message was displayed in the center of the screen that indicated whether the response was “correct” (green text) or “incorrect” (red text).

### 2.3 Bayesian rule-learning model

The rule-learning model was based on the "rational rules" model (Goodman et al. 2008) and implemented in the Python library LOTlib. This model formalizes a statistical learner that operates over the hypothesis space of Boolean propositional logic expressions (e.g. (A AND B) OR C). It implicitly defines the infinite hypothesis space of possible expressions using a grammar that permits only valid combinations of logical primitives (AND, OR, NOT) and observable perceptual features. This grammar also defines a prior probability P(H), that biases learners to prefer short expressions (H) that re-use grammar rules. The prior is combined with a likelihood, P(D|H), which quantifies how accurately a rule H predicts observed true/false labels.

We used Markov-Chain Monte-Carlo to perform inference by sampling hypotheses according to P(H|D). To predict incremental responses as the task progressed, sampling was also run incrementally for 10,000 steps at every trial, based on observed data up to that point. The top 100 hypotheses for any amount of data and concept were collected into a set that was treated as a finite hypothesis space for the purposes of efficiently computing model predictions (Piantadosi et al. 2012).

### 2.4 Reinforcement Learning models

We designed our task to study the mechanism by which humans learn using explicit rules or concepts. Nonetheless, reinforcement learning is a powerful algorithm that can perform well in most tasks, including ours. We compared different reinforcement learning (RL) models with different state space representations. Naïve RL had one feature for each stimulus (e.g. blue striped square). Feature RL had a different feature dimension for each stimulus feature (i.e., blue, striped, and square), and weights for each feature were learned independently. Exhaustive RL (designed to perform best on our task) had a state associated with each stimulus feature, each pairwise combination of features, and each triplet of features. The value of a state-action pair on trial t was determined by taking a weighted sum of each of the feature-action pairs:

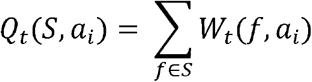

where there are two actions (a_i_) associated with the two choices (Bim or Dax). After trial feedback, the model uses RL to update each feature weight for the next trial:

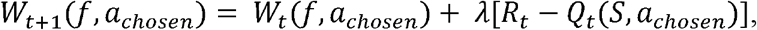

where λ is the learning rate and R_t_ is the reward earned. We departed from standard Q learning by making two opposite updates to the weight of each feature-action pair:

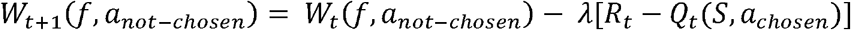

This encodes the symmetry of the task (i.e., if a stimulus is a Bim then it is not a Dax). It also improved the likelihood of the observed data without adding an extra parameter, which favored RL in model comparison. This change also made the model homologous to a standard value learning model that directly learns the probability that a stimulus is a Bim.

Each model used a softmax decision rule to map task state Q values to the probability of a given action:

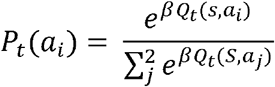

Each RL model had two free parameters: λ (learning rate) and β (inverse temperature of the softmax).

In addition, we fit a version of each RL model with a Pearce-Hall style update that augments learning rates for states with recent surprising outcomes (Pearce and Hall 1980; Li et al. 2011):

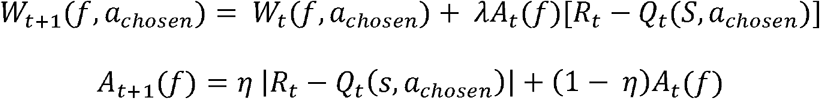

Where *A_t_(f)* is the *associability* of feature *f* at trial *t*. An additional parameter, *ŋ*, governs the degree to which associability depends on recent trials. If *ŋ* = 1, then associability is fully dependent on the previous trial, whereas if *ŋ* = 0, associability never changes from its initial value, 1, and the model reduces to standard reinforcement learning.

Finally, we fit a mixture model of RL and Bayesian rule learning that took a weighted combination of the choice probabilities from both models. The probability of the action on trial *t* was modeled as:

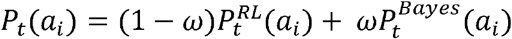

We fit models by maximizing the likelihood of observed choices for each subject, using Scipy’s minimize function with the BGFS method. For neuroimaging analyses, we calculated a single λ and β across subjects, which provides regularization that biases results away from extreme parameter settings (Daw et al. 2011).

### 2.5 Model Comparison

To compare different models, we used a leave-one-rule-out cross-validation approach (Niv et al. 2015). For each model and rule, the model was fit to the data of the remaining rules for each subject. The model, along with the maximum a posteriori parameter estimates, was then used to predict choice on each trial of the held-out rule block. We computed the average likelihood per trial of the held-out rule for each subject. This metric varies from 0 to 1, where .5 is the expected chance performance of a null model and 1 is perfect prediction. This approach allows for comparison of non-nested models with different numbers of parameters, as overfitting is naturally penalized by a reduced out-of-sample prediction accuracy.

### 2.6 fMRI acquisition

Functional images were acquired with a 3T General Electric Discovery scanner (Waukesha, WI, USA). Whole-brain BOLD weighted echo planar images were acquired in 40 oblique axial slices parallel to the AC-PC line with a 2000 ms TR (slice thickness = 3.4 mm, no gap, TE = 30 ms, flip angle = 77°, FOV = 21.8 cm, 64 × 64 matrix, interleaved). High-resolution T2-weighted fast spin-echo structural images (BRAVO) were acquired for anatomical reference (TR = 8.2 ms, TE = 3.2 ms, flip angle = 12°, slice thickness = 1.0 mm, FOV = 24 cm, 256 × 256).

### 2.7 fMRI analysis

Preprocessing and whole brain analyses were conducted with Analysis of Functional Neural Images (AFNI; Cox, 1996). Data were slice-time corrected and motion corrected. No participant included in the analyses moved more than 2 mm in any direction. Data used in whole brain analyses were spatially smoothed with a 4 mm FWHM Gaussian filter. Voxel-wise BOLD signals were converted to percent signal change.

We transformed the T2-weighted structural image to Talairach space and applied this transform to preprocessed functional images. Normalized functional images were then analyzed using a general linear model in AFNI. The model contained multiple regressors to estimate responses to each task component, which were then convolved with a two-parameter gamma variate hemodynamic response function. For surface plots, we projected the group data onto the freesurfer template brain using Pysurfer with the default settings and a 6 mm cortical smoothing kernel.

Parametric regressors derived from the Bayesian model were used to identify brain regions where activation scaled with trial-by-trial estimates of surprise, as well as the degree of change to the hypothesis space as each new exemplar was integrated. Surprise was calculated as the probability against the label assigned to the stimulus on each trial by the model. Rule updating was calculated as the KL divergence between the posterior distribution on hypotheses before and after the trial. Rule updating and surprise were moderately correlated, *r*(12) = 0.27, p = .003, but they were included in the same linear model to ensure that each captured a distinct component of neural activation. Our model included 1) surprise at feedback, 2) rule updating at feedback, and 3) rule updating from the previous trial during the subsequent cue period.

Each event in our general linear model was assumed to occur over a 2 s period (boxcar). We included three additional regressors to model mean activation during the cue, response, and feedback periods. A parametric regressor was included to control for activation that varied with reaction time during the response period. Finally, regressors of non-interest were included to account for head movement and third-order polynomial trends in BOLD signal amplitude across the scan blocks.

Maps of beta coefficients for each regressor were resampled and transformed into Talairach space. Whole-brain statistical maps were generated using one-sample t-tests at each voxel to localize brain areas with significant loadings on regressors across subjects. Whole-brain maps were thresholded at *p* < .05, cluster corrected (*p* < .005 voxel-level α with a minimum of 42 contiguous voxels, AlphaSim). Of note, we used AFNI version 16.0.06, in which AlphaSim better estimates type II error. Coordinates are reported in Talairach space with the LAS convention.

A hierarchical analysis was conducted to assess neural responses to Bayesian surprise and RL prediction error. First, activation was modeled with two regressors that encoded activation during positive and negative feedback, as well as task and nuisance variables described above. A second analysis was performed on the residual values obtained from the first regression. Two regressors were included in the second model: a parametric regressor encoding activation that scaled parametrically with surprise during feedback, and a parametric regressor encoding activation that scaled with prediction error during feedback, with prediction error derived from the best fitting (exhaustive) RL model. Because the prediction errors of the Pearce-Hall version were highly correlated with those of the standard version, r(118) = .98, and they had very similar predictive likelihoods, we opted to use the errors from the simpler RL model. To test for voxels that varied with RL prediction error, we performed a conjunction between the first-level analysis of positive > negative outcomes and the second-level parametric prediction error regressor. To test for voxels that varied with surprise, we performed a conjunction between the first-level analysis of negative > positive outcomes and the second-level parametric surprise regressor (see Results section 3.3 for the finding that surprise was greater following negative than positive outcomes). We repeated the above analysis separately for the parametric rule updating analysis to ensure that the cIFS cluster we identified did not merely respond to outcome valence. Additionally, we directly compared surprise > RPE, as well as RPE > baseline, without requiring a conjunction with outcome valence.

We analyzed outcome valence serially and in conjunction with parametric error signals for two related reasons. First, testing for a correlation with error alone is likely to identify regions that truly only have bimodal responses based on outcome. This is because outcome valence accounted for 54% of variability in prediction error variance in our data. Various effects that are not of direct interest in this study (e.g. attention, arousal, sensory differences during positive versus negative feedback) will differ by outcome and we wished to diminish the influence such spurious factors in our results. Second, factoring out outcome valence, which would eliminate such spurious results, dismisses a critical component of RL. An RL model absent outcome valence information cannot learn to distinguish actions and will respond randomly, while a RL algorithm that has access only to outcome valence, but does not remember recent trials, will implement a win-stay/lose-shift policy. Therefore, it is necessary to examine outcome valence and parametric error signals together (i.e., in a conjunction analysis) in order to make strong conclusions about the relationship of the BOLD response to prediction error or surprise.

We conducted an ROI-based analysis of the evoked response in the striatum. Based on the results of the rule-updating analyses, in which cIFS correlated with rule-updating and was functionally connected with the striatum, we hypothesized that surprise might be represented in striatal regions that interact with lateral PFC (Haber and Knutson 2009) . We used an “executive caudate” ROI taken from a 3-way subdivision of striatum based on diffusion tractography imaging estimated connectivity with cortex crossed with a caudate anatomical ROI (Tziortzi et al. 2013). We extracted preprocessed data from this ROI and regressed out 1) head motion, 2) third-order polynomial trends, and 3) cue, response, and reaction-time related activation, as in the main GLM. For statistical analysis, we averaged the response between 6-10 seconds after feedback. We analyzed data using a mixed linear model with a random slope for each subject as implemented in LMER in R.

## 3 RESULTS

### 3.1 Behavior

We collected fMRI data while participants completed six 20-trial blocks of a rule-learning task (Figure 1a). In each trial, participants were shown an image and were instructed to classify it as belonging to one of two possible categories (“Dax” or “Bim”) based on perceptual features. Category membership was determined based on logical rules like “stimuli that are either blue or square are Bims”. Learning proceeded rapidly, generally reaching an initial asymptote within the first five trials (Figure 1C). For all but the simplest rules, accuracy diminished on later trials, evidenced by accuracy curve “spikes.” These accuracy drops occurred on trials where new, highly informative evidence was presented that required updating from a simpler to a more complex rule.

**Figure 1:**
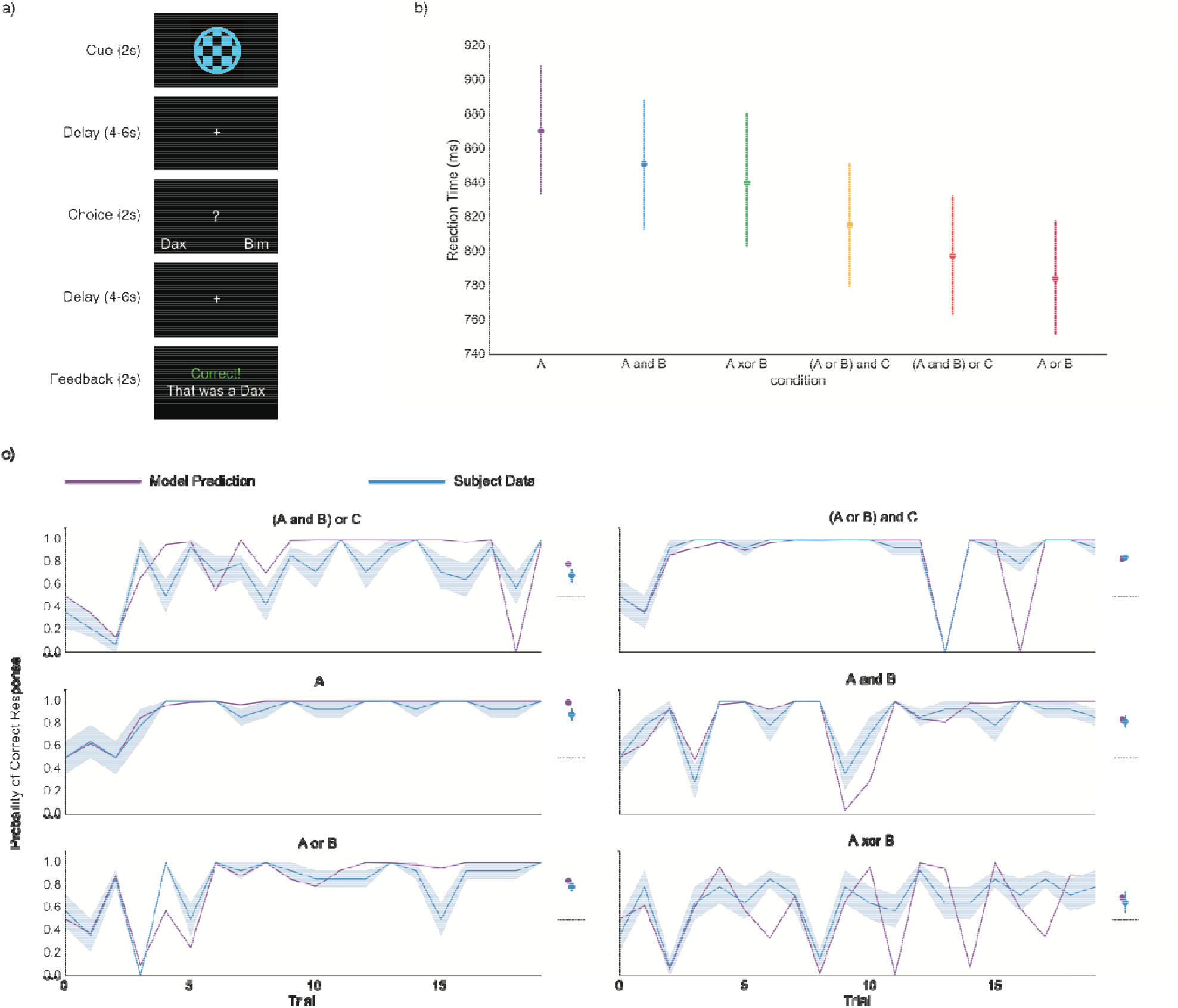
Rule Learning Task and Behavior. a) Participants completed six 20-trial blocks of a rule-learning task. Trials were divided into three phases: cue, response and feedback, each separated by a random 4-6 s delay. During the cue phase (2 s), the stimulus to be categorized was presented in the center of the screen. During the response phase (2 s), a question mark was presented in the center of the screen, prompting participants to press a button to respond. During the feedback phase (2 s), a message was displayed indicating whether the response was correct. b) Average reaction times for each of the rule blocks, ordered by mean reaction time. Although there was heterogeneity in reaction time between rules, only the difference between A and (A and B) was significant when correcting for multiple comparisons. c) Left panels show mean participant accuracy and Bayesian rule learning model predictions, without any parameter fitting, for each rule. To the right is the average performance, collapsed across trials, referenced against the performance of an optimal version of the Bayesian model for each rule. The dotted line represents chance performance. Participants learned to respond well above chance and remarkably close to optimal performance for all rules. All error bars represent bootstrapped estimates of the standard error of the mean across subjects.

All participants included in analyses performed above chance in categorizing stimuli. Accuracy was significantly greater than chance for all rules (Figure 1C, A: *t*(13) = 14.7, *p* < .001; (A and B) or C: *t*(13) = 7.1, *p* < .001; A or B: *t*(13) = 16.0, *p* < .001; A and B: *t*(13) = 12.9, *p* < .001; (A or B) and C: *t*(13) = 33.7, *p* < .001; A XOR B: *t*(13) = 3.25, p = .038; all ps Bonferroni corrected). There was an effect of condition on accuracy (within-subjects ANOVA with Greenhouse-Geisser spherecity correction, F(3, 38.6) = 10.1, *p* < .001 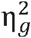 = 0.40, Tables S2). There was additionally an effect of condition on reaction times, F(2.7, 26.8) = 3.1, p = .03, 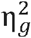 = 0.02 (Table S3).

Accuracy is not the ideal measure of learning, as subjects are expected to sometimes answer incorrectly. Figure 1C shows on the right panels average accuracy compared to the accuracy of an ideal Bayesian model that deterministically selects the rule with highest likelihood. Although the Bayesian model outperformed subjects, z = 4.5, *p* < .001, the average difference in accuracy between subjects and the model was only 5.2% . Therefore, subject behavior reflected near-optimal performance given the demands of the task.

### 3.2 Comparing Bayesian and RL Models

Our task was designed to elicit an explicit learning strategy that relied on the generation and testing of abstract rules. Nonetheless, reinforcement learning is a powerful learning algorithm that can perform well on many tasks, including ours. In addition, there are well-established cortico-striatal circuits for stimulus-response and feature-response learning that could contribute to performance on our task (Niv 2009). To strengthen our claim that subjects adopted a rule-learning strategy, we used computational modeling of behavior to assess the extent to which subjects employed rule-learning rather than model-free RL.

A Bayesian model of rule learning predicted 78.5% of the variance in subjects’ choices without any parameter fitting. This accuracy was well-above chance, *t*(13) = 29.9, *p* < .001, replicating previous work employing similar models (Figure 1B; (Goodman et al. 2008; Piantadosi 2011). In spite of a large effect of condition on model-fit, F(2.5, 32) = 27.7, *p* < .001, 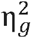 = 0.62, Figure 2B, Table S4, predictive accuracy was well-above chance for all rules, A: *t*(13) = 22.5, *p* < .001; (A and B) or C: *t*(13) = 8.3, *p* < .001; A or B: *t*(13) = 23.7, *p* < .001; A and B: *t*(13) = 20.8, *p* < .001; (A or B) and C: *t*(13) = 34.5, *p* < .001; A XOR B: *t*(13) = 4.2, p = .006; all ps Bonferroni corrected.

**Figure 2:**
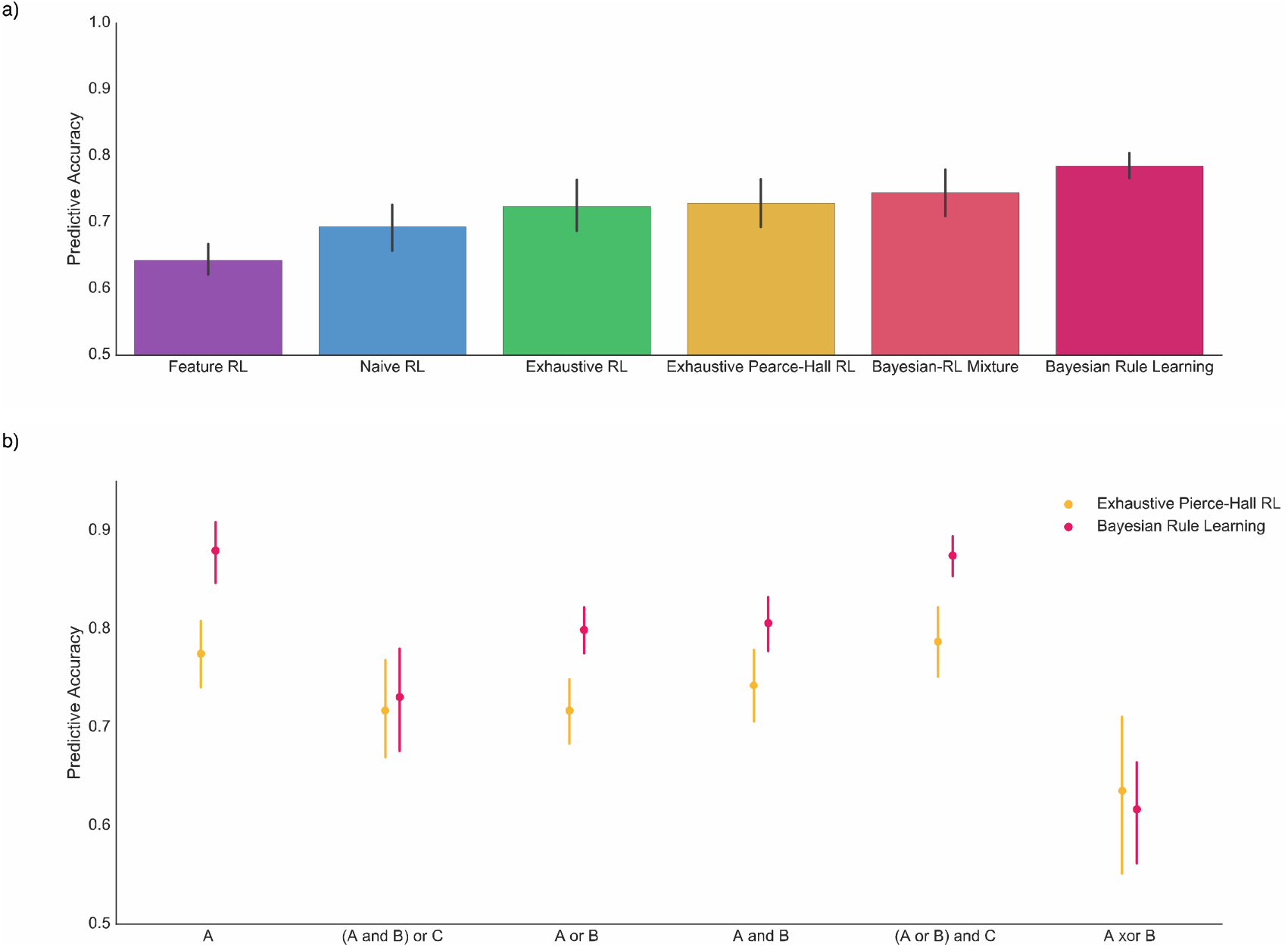
Bayesian Rule Learning outperforms Reinforcement Learning models. a) Bayesian Rule Learning outperforms standard reinforcement learning models that learn about either stimuli (Naïve) or features (Feature) in predicting subject behavior. The model also outperforms an RL model with an exhaustive state space (features, stimuli, and pairwise combinations of features) as well as an augmented version of this model with a Pierce-Hall update. Finally, the Bayesian model outperforms a mixture model that combines both Bayesian and RL predictions. b) Rule-by-rule comparison of the predictive accuracy of the Bayesian model and the best performing, exhaustive Pearce-Hall RL model. Despite significant heterogeneity, the Bayesian model outperforms the RL model for most rules, whereas the RL model does not significantly outperform the Bayesian model on any rule.

We analyzed RL models that are 1) commonly used in the literature to describe behavior and 2) are based on the established circuits between sensory cortex and the striatum that support model-free learning (Niv 2009). We contrasted the Bayesian model fits with fits produced by five RL models of varying complexity.

The first model, naïve RL, learns independently about each stimulus. Since each stimulus repeats 2.5 times, on average, this model performs quite well. In addition, this model has described behavior well in related tasks (Niv et al. 2015). Because our task is noiseless, it also describes an approach to the task based on episodic memory.

The second model, feature RL, learns about each feature (e.g., blue, circular, and striped) independently. This model reflects the idea that the decomposition of stimuli into constituent features in sensory cortex can support cached-value learning based on those features (O’Reilly and Rudy 2001). Feature RL could be expected to learn some rules well, {A, A or B}, but should struggle with rules that involve conjunctions of features {A and B}.

We constructed the third and fourth RL models to be ideally suited to learning the rules in our specific task. This third RL model, exhaustive RL, learns weights for all stimuli (unique combinations of shape, color, texture), each individual feature, and each pairwise combination of features. For the fourth model, we fit a version of exhaustive RL that incorporates an attention-like mechanism that updates features more if they have been associated with surprising outcomes in the past (exhaustive Pearce Hall; (Li et al. 2011).

Finally, for the fifth model, we created a mixture model that takes a weighted average of the predictions of the Bayesian model and the Pearce-Hall RL model with the exhausted state space. This model directly tests whether RL provides any additional explanatory power over the Bayesian rule model.

To compare the ability of each of the models to account for subjects’ behavior, we fit each model on 5 of the 6 rules and contrasted the likelihood of the predicted held-out rule data. This predictive likelihood approach balances model fit and complexity because over-complex models will over-fit the training data and generalize poorly.

The Bayesian rule model predicted unseen behavior better than standard RL models (feature: *t*(13) = 8.1, *p* < .001; naïve: *t*(13) = 3.8, *p* = .002). As expected, both exhaustive RL models outperformed the best standard RL model (naïve) in accounting for behavior (exhaustive: *t*(13) = 3.3, *p* = .005, exhaustive Pearce-Hall: *t*(13) = 4.2, *p* = .001). Further, the exhaustive Pearce-Hall model outperformed the exhaustive model, *t*(13) = 2.4, *p* = .034, replicating prior work (Li et al. 2011). Strikingly, the Bayesian rule learning model outperformed even the exhaustive Pearce-Hall model, *t*(13) = 5.1, *p* < .001. In summary, the Bayesian model predicted behavior better than all tested RL models.

We next examined whether there was variability between rules in the relative abilities of the models to explain behavior (Figure 3b). Although there was an effect of rule on the difference between Bayesian and RL model fits, F(3.4, 44.1) = 16.3 *p* < .001, 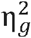 = 0.38, Table S5, the best RL model outperformed the Bayesian model numerically, but non-significantly, only for XOR, *t*(13) = −1.1, *p* > .3.

**Figure 3:**
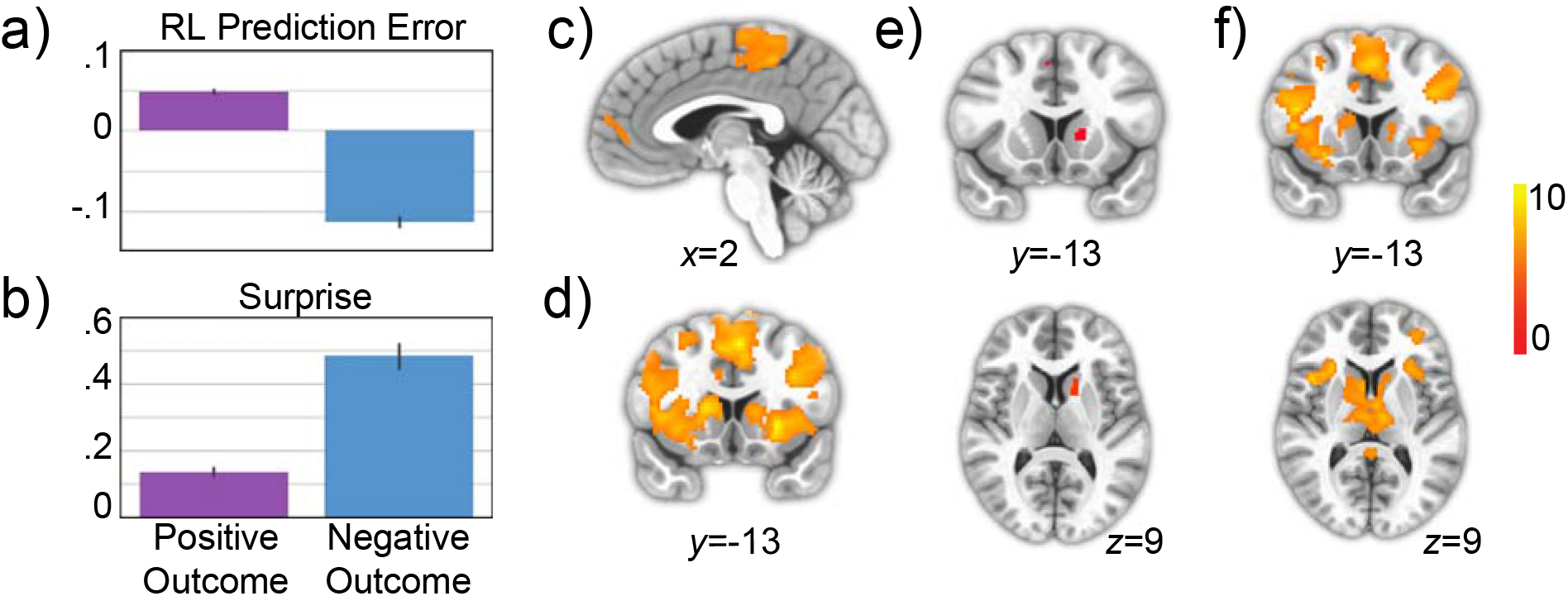
Striatum represents Bayesian surprise, not reinforcement learning prediction error a) Mean prediction error from the best fitting RL model, sorted by whether outcome was positive or negative. b) Mean surprise from Bayesian rule learning model, sorted by whether outcome was positive or negative. c) Whole brain corrected results for the contrast of positive > negative outcomes. There were no significant voxels in the striatum for this contrast. d) Whole brain corrected results for the contrast of negative > positive outcomes. e) Results of a conjunction analysis displaying voxels that are significantly active for both negative > positive outcomes and the parametric effect of surprise. Both contrasts were corrected for multiple comparisons across the whole brain before being entered into the conjunction analysis. f) Whole brain corrected results for the contrast of parametric surprise > parametric prediction error, without the effect of outcome partialed out.

The fifth model, a mixture that allow for combined input from the Pearce-Hall and Bayesian models, provides the definitive test of the descriptive quality of the Bayesian model. If RL provides any addition descriptive information over the Bayesian model alone, then the mixture model should provide superior predictions of behavior. Although this model predicted behavior better than the best RL model, *t*(13) = 3.0, p = .011, indicating that the Bayesian model provides unique information, it was worse at predicting behavior than the Bayesian model, *t*(13) = 4.2, *p* < .001. This result indicates that mixing RL predictions with Bayesian predictions results in overfitting without any additional explanatory power over the Bayesian model alone (see supplemental materials for further discussion of modeling). We conclude that our task manipulation successfully biased subjects towards a rule learning, rather than a reinforcement learning, strategy.

### 3.3 Striatum activation does not reflect reinforcement learning prediction errors

Our behavioral analysis suggested that our task design successfully biased participants towards rule-learning. We next tested whether the striatum reflects prediction errors, which would underlie incremental stimulus response learning, or Bayesian “surprise”, which reflects beliefs about abstract rules. The learning signal used by RL algorithms is the prediction error, which is the difference between the prediction strength of the chosen action and an indicator function on whether the action was correct. In our case,

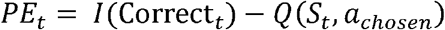

The analogous signal in our Bayesian rule-learning model, which we call surprise, is equal to the difference between the strength of evidence for a category and the actual category. If on trial *t* a given stimulus *S_t_* is a Bim (0_t_ = Bim), then the surprise is given by

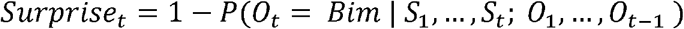

Although RL prediction errors and “surprise” are generally correlated, they diverge in an important way. RL prediction error is signed and therefore will almost always be greater for correct than for incorrect responses. Although “unsigned prediction errors” exist in other modeling frameworks such as predictive coding, and are represented in different neural circuits; temporal difference errors are signed and strongly associated with VTA dopamine neurons and striatal BOLD responses. The signed error is what endows the model with the ability to prefer states and actions that lead to rewards over those that do not. In contrast, surprise will be larger for incorrect than correct outcomes because incorrect predictions are more surprising in our task (see below). This implies that there would be trials with large positive surprise and large negative prediction errors.

Surprise was larger for negative outcomes (t = 21.8, p = 3.7 ×10^−93^, Figure 3b), and outcome valence alone accounted for 22.1% of the variance in surprise. Conversely, RL prediction errors extracted from the best-fitting exaustive model were larger for positive outcomes (t = 47.2, p = 2.3 ×10^−310^, Figure 3a) and outcome valence alone accounted for 57.1% of the variance. The effect of outcome valence on these learning signals was large and in opposite directions. These effects are useful because they allow for a simple and non-parametric test of RPE: a region that does not show a larger response to positive than negative outcomes cannot have a positive relationship with RPE (see Methods for discussion). We therefore predicted that, if the striatum reflects surprise in our task, then mean striatal blood oxygen level dependent (BOLD) activation should be larger for negative compared to positive outcomes. This is a novel prediction given the extensive body of work showing larger striatal responses to positive than negative outcomes (Delgado 2007). We emphasize that this prediction is specific to our particular task in which negative feedback is generally more informative than positive feedback.

An analysis of negative versus positive outcomes yielded a large cluster of voxels that encompassed most of the dorsal striatum as well as part of the ventral striatum (Figure 3d). The reverse contrast (positive vs. negative outcome) yielded regions in the dorsomedial prefrontal cortex (dmPFC) and posterior paracentral lobule (pPCL; corrected *p* < .05, Figure 3c), but no striatal regions were found, even at liberal thresholds (*p* < 0.01, uncorrected). Together, these results are incompatible with a positive relationship between striatal BOLD and RL prediction error.

### 3.4 Striatum activation varies with Bayesian surprise

We next turned to a model-based analysis to probe whether trial-by-trial fluctuations in surprise account for striatal activation. We observed a region of the caudate that was sensitive to surprise, in addition to regions in supplementary motor area (SMA; *p* < .05 corrected, Figure 3e). Conversely, we did not observe any significant RL prediction error associated activations at our whole brain threshold, and did not observe any in the striatum even at a lenient threshold (*p* < 0.005, uncorrected). Further, we did not observe any striatal activation for an RL prediction error signal even without masking by outcome valence (*p* < 0.005, uncorrected). Because the conjunction criterion established that the striatal surprise response was not due to outcome alone, we next formally compared surprise and prediction error regressors without projecting out variance due to outcome. This analysis rules out any potential impact of removing the outcome variance associated with surprise or prediction error. We observed a robust response in the striatum for the contrast of surprise > prediction error (*p* < .05 corrected, Figure 3f), but did not observe any activation for the reverse contrast in the striatum, even at a lenient threshold (*p* < 0.005 uncorrected).

In order to investigate striatal activation in more detail, we examined the evoked hemodynamic response to feedback in our executive caudate ROI. The evoked response to negative feedback was larger than to positive feedback, z = 5.63, *p* < .001 (Figure 5a). We next modeled the evoked response as a function of surprise, prediction error, and each signals’ interaction with rule condition (deviation coding). Surprise was significantly related to the evoked response, z = 4.1, *p* < .001 (Figures 5b, S4), whereas RPE was not, z = −1.46, p = .145 (Figure 5c). There were no significant interactions with rule condition for either signal (all *p* > .1, Figures 5d, S5). Together with the results of the whole-brain analyses, we conclude that in our rule-learning task, striatal activation reflects Bayesian surprise rather than RL prediction error. This result is consistent with our behavioral analysis that concluded that subjects reason over rules rather than incrementally acquired stimulus-response contingencies.

**Figure 4:**
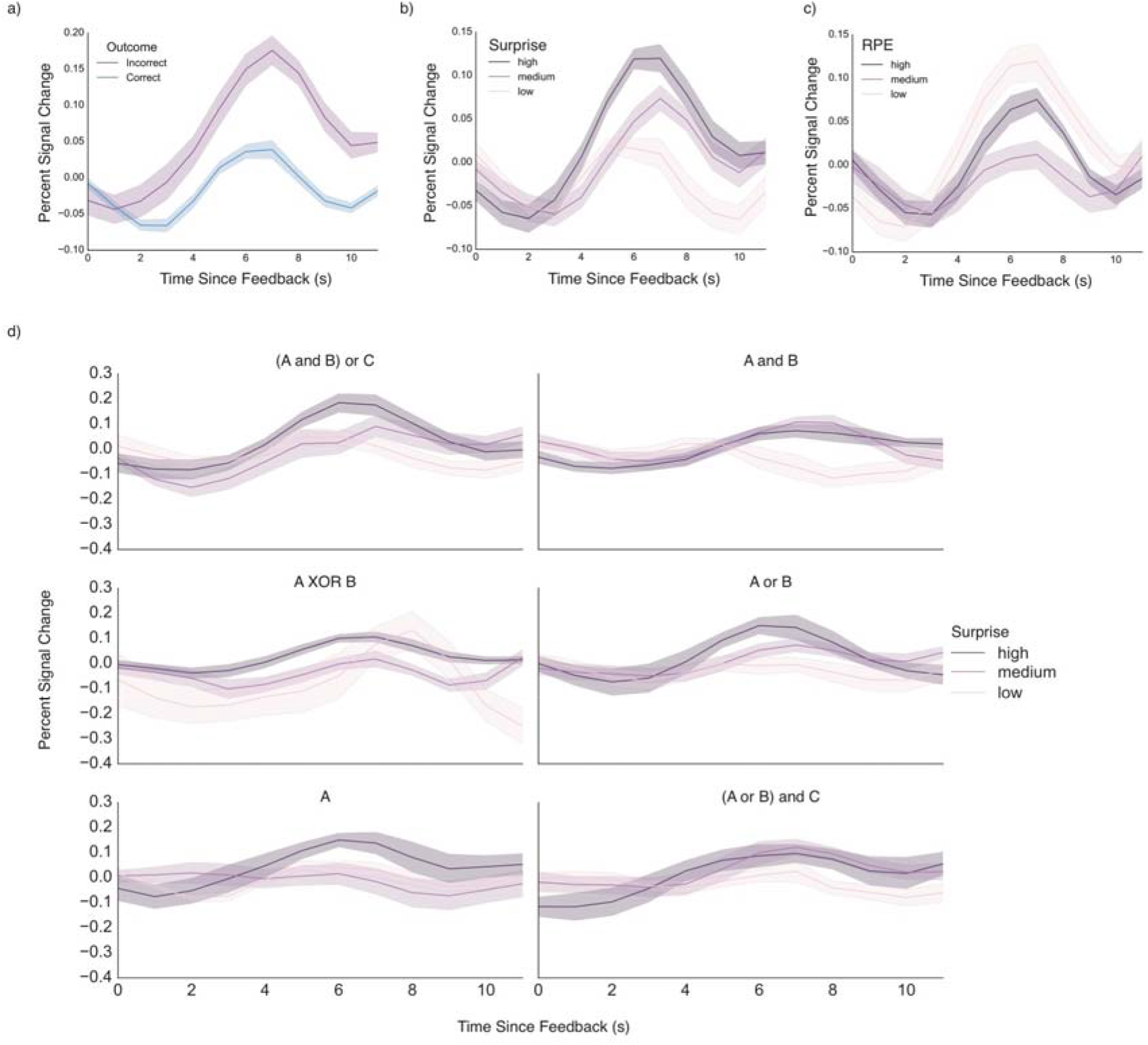
Analysis of feedback response from a caudate ROI defined based on its connectivity to executive cortical areas. a) Responses to negative outcomes were greater than to positive outcomes. b) A monotonic relationship existed between Bayesian surprise and response amplitude, with greater response for highest surprise. C) By contrast, no monotonic relationship was evident between striatal response and reward prediction error. D) Striatal activity is plotted as a function of surprise for each rule learned in the task. Despite some heterogeneity, the striatum generally increases its response as a function of surprise across rules. Error bars represent bootstrapped estimates of the standard error of the mean.

**Figure 5:**
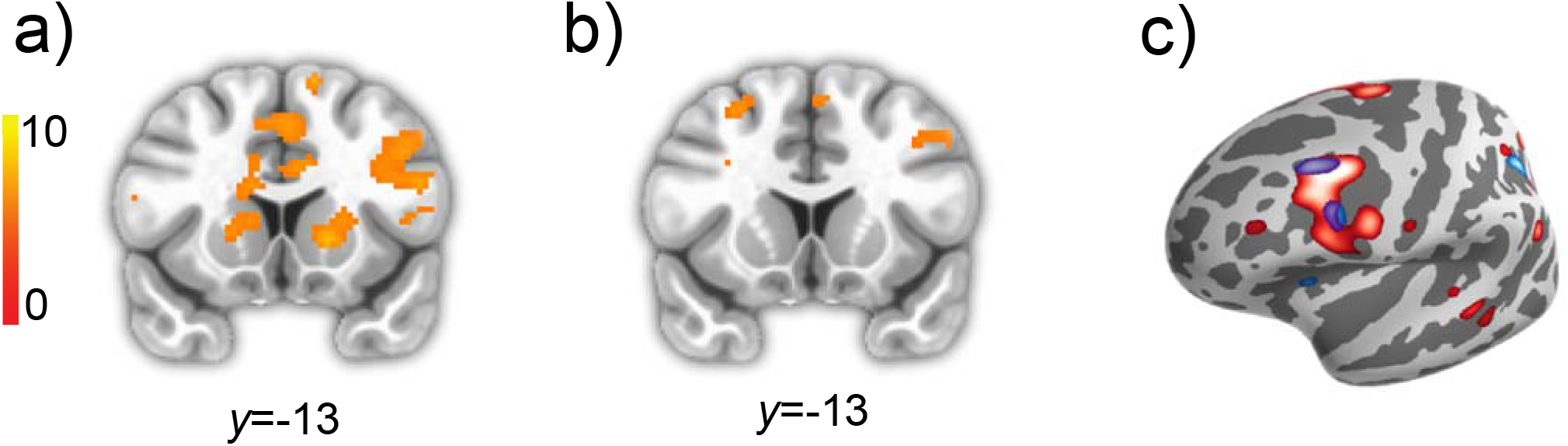
Rule Updating. a) Rule updating during the feedback period in the striatum and left cIFS b) Rule updating during the subsequent cue period in the left cIFS. c) Projections of a and c onto the cortical surface. Red corresponds to rule updating during the feedback period, blue corresponds to rule updating during the subsequent cue period.

### 3.6 Rule updating

Our analyses show that the Bayesian model described striatal activation better than RL prediction errors in a rule learning task. However, the striatum also tracks RPE in reinforcement learning tasks (Rutledge et al. 2010). These observations can be reconciled under a model where striatal neurons represent values and these values change in response to errors. In tasks for which RL describes the learning process, errors should come from a RL-like algorithm, and in a rule-learning task like ours they should reflect errors in beliefs about rules. This model suggests that apparent BOLD error signals in the striatum can be better characterized as changes in value signals. However, it is very difficult to distinguish value updating and reward prediction error in existing work that uses RL models of BOLD data because the value update and reward prediction error are proportional (their ratio is the learning rate). Conversely, our Bayesian model allows us to separately examine representation of surprise and rule updating.

As a further corollary to this model, we expected that rule updating would involve both the striatum and the caudal inferior frontal sulcus (cIFS), because of the known role of the cIFS in feature-based rule maintenance and execution (Koechlin et al. 2003; Badre and D'Esposito 2007; 2009; Waskom et al. 2014). Specifically, we hypothesized that if the cIFS maintains and executes the feature-based rule governing behavior, then neural activity in this region should change when the rule governing behavior was subject to change. We used trial-to-trial KL divergence in rule likelihoods as our measure of rule updating, and expected that this measure would predict shifts in internal representation of rule likelihoods. In a whole-brain regression analysis, we identified brain regions correlated with rule updating at the time of feedback in the bilateral cIFS, intra-parietal lobule (IPL), fusiform gyrus, and portions of the dorsal caudate (Figure 5).

Although rule updating occurs during feedback, it is likely that some updating of rule-response contingencies happens during subsequent cue periods. Evidence for this comes from task-switching paradigms, in which subjects incur a residual switch cost even when they have ample time to prepare for the new task (Sohn et al. 2000; Monsell 2003). This may be taken as evidence that people do fully encode a new rule until it must actually be implemented. We observed activation patterns corresponding to rule updating during the cue period of the subsequent trial in the cIFS and fusiform gyrus bilaterally (Figure 5). A conjunction analysis of rule updating during feedback and the next cue confirms that these areas track rule updating during both time periods (Figure 5; (Nichols et al. 2005), and follow-up analyses established that these results were not driven by outcome effects (Figure S1). Our findings are consistent with a model in which the cIFS maintains and updates the rules governing behavior, while the striatum maintains and updates the values of the dominant and competing rules.

## 4.1 DISCUSSION

The brain is adept at building stimulus-response relationships, but both humans (Goodman et al. 2008) and nonhuman primates (Costa et al. 2015) exploit structured relationships in order to learn efficiently. We designed a task in which participants are biased towards reasoning about explicit rules to categorize stimuli, rather than relying on the gradual build-up of stimulus-response contingencies. We found that the striatal feedback response tracks Bayesian surprise, rather than reward prediction error, in this learning context. We additionally found that both the striatum and cIFS track changes in beliefs about rules. Together, these results suggest that cortico-striatal interactions support learning about rule-based structured relationships.

Our behavioral analysis confirmed that our task design elicited behavior that was consistent with a Bayesian optimal rule learning strategy. We exploited a difference between the feedback signals generated from RL models (reward prediction error) and Bayesian rule learning (surprise). RL prediction error is a signed learning signal that is larger for positive than negative feedback; in contrast, surprise is larger for negative feedback in our task. Negative feedback generated a stronger response in the striatum, which is inconsistent with an RL prediction error account of the striatal feedback response in our task. Further, striatal BOLD responses tracked a parametric measure of

Bayesian surprise and did not track RPE. Finally, surprise accounted for striatal responses better than prediction error. Together, these results indicate that the striatal feedback response reflects Bayesian surprise in task conditions where behavior is governed by explicit reasoning about and error-driven updating of abstract rules.

The association of the striatal feedback response with reward prediction error saturates the human cognitive neuroscience literature. Several studies have hypothesized that reward prediction error representation is the primary function of striatum during learning (Hare et al. 2008; Daw et al. 2011; Garrison et al. 2013). Indeed, in reinforcement learning tasks, the striatal feedback response clearly tracks reward prediction error (Rutledge et al. 2010). These differences across paradigms can be accounted for if the striatal feedback response reflects the change in values encoded by striatal neurons in response to new information. This more general account of the striatal feedback response is bolstered by several experimental observations. First, striatal response to negative feedback increases if the feedback is less predictable (Lempert and Tricomi 2015). Second, the striatum responds more to negative feedback that indicates a set-shift in the Wisconsin card sorting task (Monchi et al. 2001). Finally, cyclic voltammetry measurements in rodents indicate that striatal dopamine appears to track a value, rather than prediction error (Hamid et al. 2015), although striatal dopamine has a complex relationship with striatal BOLD (Lohrenz et al. 2016). To build upon these observations, we leveraged model-based fMRI and a novel task to examine the striatal feedback response under conditions where learning does not depend on the incremental adjustment of stimulus-response contingencies. Future work measuring the response properties of striatal neurons in humans should directly probe the information carried by these neurons during rule learning.

To bolster our claim that the striatal feedback response is linked to its role in updating internal value representations, we showed that the striatal BOLD covaried with a parametric measure of rule updating. Our design afforded us a unique opportunity to separately examine error and updating. In standard reinforcement learning these quantities are directly related so that they may not be distinguished analytically. Our analysis identified a region in prefrontal cortex called the cIFS, which is involved in the maintenance and execution of rules that depend on the features of visual stimuli (Koechlin et al. 2003; Badre and D'Esposito 2007). Our observation that cIFS activation scaled with rule updating supports a model in which this area selects and executes rules based on rule value representations in the striatum. Future work will investigate the chain of sensory processing that connects visual features to abstract decision rules in cortex. The hippocampus may play a critical role in this process (Mack et al. 2016).

Our results are related to the findings that striatal responses to feedback reflect both model-based and model-free RPE (Daw et al. 2011), whereas prediction errors related to knowledge of the environment are reflected in lateral prefrontal cortex (Glascher et al. 2010). Our finding that rule updating correlates with cIFS activation is consistent with this latter finding. However, we find no evidence of an RPE signal in the striatum. We believe the discrepancy is derives from the fact that the probabilistic and evolving nature of the two-step task favors an RL system. This system stores stimulus-value associations in striatal synapses, draws on these associations to make predictions, and updates them in response to reward prediction errors. In our task, the values of rule representations may be stored in a separate population of striatal synapses, drawn on to make predictions, and updated in response to surprising outcomes. The nature of the stored associations and the way they are updated may differ between tasks, but the tasks are similar to the extent that knowledge represented in cortex influences predictions about reward. Future work could examine model-based learning as a set of mechanisms by which cortical representations interact with stored associations in the striatum to drive behavior.

We did not use explicit rewards, such as money, in our task, which differs from some RL experiments. However, the striatum is consistently sensitive to feedback in a manner that is similar to its response to explicit rewards (Elliott et al. 1997; Seger and Cincotta 2005; Tricomi et al. 2006; Marco-Pallarés et al. 2007; Dobryakova and Tricomi 2013; Swanson and Tricomi 2014; Lempert and Tricomi 2015). Also, the striatum has been shown to respond to internally generated reward prediction errors used in hierarchical RL (Ribas-Fernandes et al. 2011; Diuk et al. 2013; Iglesias et al. 2013). Both empirical and theoretical work suggests that the brain’s learning system uses surrogate rewards to learn in the absence of monetary reward receipt.

This study tested the simple idea that the role of the striatum during learning is to compute the value of potential behavioral policies and update them in response to new information. We delineated the functional neuroanatomy underlying rule-based learning and in the process ruled out a RL account of striatal activation in deterministic category learning. Our results suggest that value updates in cortico-striatal connections facilitate rule-based learning.

## 5.2 ACKNOWLEDGEMENTS

The authors would like to thank G. Elliott Wimmer, Vishnu Murty and Kim D’Ardenne for helpful feedback.

